# Fecundity determines the outcome of founding queen associations in ants

**DOI:** 10.1101/2020.11.20.391359

**Authors:** Eva-Maria Teggers, Falk Deegener, Romain Libbrecht

**Affiliations:** Institute of Organismic and Molecular Evolution, Johannes Gutenberg University of Mainz, Germany

**Keywords:** Pleometrosis, Social insects, Cooperation

## Abstract

Animal cooperation evolved because of its benefits to the cooperators. Pleometrosis in ants - the cooperation of queens to found a colony - benefits colony growth, but also incurs costs for some of the cooperators because only one queen usually survives the association. While several traits are associated with queen survival, they tend to be confounded and it is unclear which factor specifically determines the outcome of pleometrosis. In this study, we used the ant *Lasius niger* to monitor offspring production in colonies founded by one or two queens. Then, we experimentally paired queens that differed in fecundity but not in size, and *vice versa*, to disentangle the effect of these factors on queen survival. Finally, we investigated how fecundity and size differed between queens depending on whether they were chosen as pleometrotic partners. Our results indicate that pleometrosis increased and accelerated worker production via a nutritional boost to the larvae. The most fecund queens more frequently survived the associations, even when controlling for size and worker parentage, and queens selected as pleometrotic partners were less fecund. Our results are consistent with fecundity being central to the onset and outcome of pleometrosis, a typical case of cooperation among unrelated animals.

## Introduction

While animal cooperation typically occurs among related individuals (Hamilton, 1964), there are cases of unrelated cooperators in some mammal, bird and insect species (Bernasconi & Strassmann, 1999). These have raised interesting questions on the origin, maintenance and benefits of cooperation among unrelated animals. Such questions include how the internal (e.g., physiology) and external (e.g., environment) conditions affect the decision to behave cooperatively, the benefits of cooperation, and its outcome. While social insects are typical model systems for kin-based cooperation, the early stage of the colony life of some ant species provides a classic example of cooperation among unrelated partners (Bernasconi & Strassmann, 1999).

Pleometrosis in ants is the foundation of colonies by two or more cooperating newly-mated queens that settle in the same nest site after a nuptial flight (Bartz & Hölldobler, 1982; Hölldobler & Wilson, 1977; Sommer & Hölldobler, 1995; Wheeler, 1917). In some cases, pleometrosis results in mature multiple-queen colonies, as in pleometrotic populations of harvester ants (Johnson, 2004; Overson et al., 2016). However, in most species, pleometrotic associations are only transitory: the queens stay together to produce eggs, but upon worker emergence, queens engage in fights (sometimes initiated and/or joined by the workers) until a single queen survives (Bartz & Hölldobler, 1982). Pleometrosis results in an earlier production of larger groups of workers compared to solitary foundation, which may be beneficial in dense populations where young colonies compete for food and brood raiding can be common (Deslippe & Savolainen, 1995; Johnson, 2004; Madsen & Offenberg, 2017; Rissing & Pollock, 1987; Sommer & Hölldobler, 1995; Trunzer et al., 1998). How pleometrosis benefits colony growth remains unclear (Brütsch et al., 2017; Pull et al., 2013).

The queens that do not survive the pleometrotic associations have no fitness benefits, as they die before producing sexuals (the fertile individuals that are only produced in mature colonies). Thus, understanding the factors that determine which queens survive the associations would allow researchers to draw and test hypotheses on whether or not queens with given characteristics should decide to cooperate with other queens, as well as understand previous reports of such queen decisions (Nonacs, 1992; Sommer & Hölldobler, 1995).

Several determinants of queen survival have been identified. Winning queens tend to be larger (Aron et al., 2009; Bernasconi & Keller, 1996; Bernasconi & Keller, 1998; Nonacs, 1990), heavier (Nonacs, 1990, 1992; but see Sommer & Hölldobler, 1995), to spend more time on the brood pile (Balas & Adams, 1996; Holman et al., 2010), and to lose less weight during colony foundation (Balas & Adams, 1996; Bernasconi & Keller, 1996). However, these reports may have been subject to confounding effects because the factors tested are usually interconnected. For example, size and mass may correlate with fecundity (Aron et al., 2009), which in turn influences worker production and therefore worker parentage in the offspring. One strategy to control for such confounded factors is to experimentally pair queens that differ in one factor but not in others, thus allowing the identification of factors that affect queen survival irrespective of others.

In this study, we used the black garden ant *Lasius niger* to study cooperation among unrelated individuals in pleometrotic associations. We first quantified the benefits of pleometrosis in terms of brood production and development. Then we tested whether fecundity and size, two factors that may be associated with queen survival (Aron et al., 2009; Balas & Adams, 1996; Bernasconi & Keller, 1996; Nonacs, 1990) were correlated. We investigated the effect of fecundity and size on queen survival in experimental pleometrotic associations that differ in one factor but not the other, while controlling for worker parentage. Finally, we compared fecundity and size between queens that were selected as pleometrotic partners and queens that were not.

## Methods

### Sample collection

We used *L. niger* as study organism because queens in this species commonly found new colonies in pleometrotic associations that consist of typically unrelated individuals (Sommer & Hölldobler, 1995). We collected 276 *L. niger* founding queens in Mainz, Germany right after nuptial flights on June 26^th^ 2017 (cohort 1, 138 queens), July 9^th^ 2017 (cohort 2, 138 queens) and July 4^th^ 2020 (cohort 3, 135 queens). The queens were housed in glass test tubes (10*1.2cm) with water blocked by a cotton ball, and then kept in the dark at 21°C.

### Experimental designs

We randomly selected 45 queens from cohort 2 to investigate the benefits of pleometrosis. Half the test tubes contained a single queen (n = 15), and the other half contained two queens (n = 15). Then we recorded the number of eggs, larvae, pupae and workers three times a week for 53 days. Upon emergence of the first workers, we placed the tubes in larger plastic boxes (11*15*3cm) and fed the colonies once a week with a mixture of honey, eggs, agar and water (Bhatkar & Whitcomb, 1970). Eight queens died over the course of the experiment: Three queens in the treatment with one queen died 4, 11 and 16 days after set-up, and were removed from the analyses. Five queens in the treatment with two queens died 37, 42 (2) and 49 (2) days after set-up. In each of those five cases, one queen of the two queens survived. These tubes were kept in the analyses because queen execution is expected in pleometrotic associations.

We used 231 queens (138 from cohort 1, 93 from cohort 2) to investigate the effect of size and fecundity on queen survival. On day 17 after the nuptial flights, we recorded the number of brood items (eggs and larvae) in each tube, and used this measure as a proxy for fecundity, thus assuming that egg production changes similarly over time across queens. On days 19 and 20 after the nuptial flights, we used a Leica stereomicroscope to measure the thorax length of the founding queens, and used it as a proxy for size in all subsequent analyses.

To test how difference in fecundity between queens affects the outcome of the pleometrotic association, we experimentally paired 70 paint-marked queens (35 pairs) that differed in the number of brood pieces (one sample t-test, t = −15.1, df = 34, p < 0.0001), but not in thorax length (one sample t-test, t = 0.59, df = 34, p = 0.56). To do so, we paired queens that had a high number of brood items (top 25% of the distribution) with queens that had a low number of brood items (bottom 25% of the distribution), while making sure that they had a medium size (both in the middle 50% of the distribution). We used a similar strategy to investigate how size difference between queens affects their survival, and paired 66 paint-marked queens (33 pairs) that differed in size (one sample t-test, t = −18.15, df = 32, p < 0.0001) but not in fecundity (one sample t-test, t = −0.36, df = 32, p = 0.72). Before being paired, each queen was marked with a color dot on the thorax using a toothpick dipped into Edding marker paint. To prevent worker parentage from being confounded with queen fecundity, we did not provide the queen pairs with their own brood, but with unrelated pupae collected in field colonies around Mainz, Germany.

We used 135 queens (cohort 3) to investigate how fecundity and size correlate with the likelihood of being chosen as a pleomeotric partner. To do so, we first performed choice test experiments on the three days following the nuptial flight. We used test arenas that consisted of two plastic tubes (length: 4 cm) covered with red foil, and connected to a plastic petri dish (diameter: 5 cm). We tethered a queen to each tube by attaching one end of a metal wire (length: ca. 1 cm; diameter: 0.02 mm) to its petiole and the other end to the bottom of the tube. We then introduced a choosing queen in the central petri dish, which could move freely into both tubes, and interact with the queens attached to the bottom of each tube. This set-up provided the choosing queen with two possible pleometrotic partners. We used Sony FDR-AX33 video cameras to record the arenas for 12 hours after introduction of the choosing queen. We tested 45 groups of three queens. For each test, we randomly selected one of the three queens to be the choosing queen. We used a set of three criteria to consider that a choosing queen had made a choice: (i) it should be observed inside one of the two tubes at the end of the video (12 hours after its introduction to the arena), (ii) it should not have left the tube for at least one hour before the end of the video, and (iii) it should be observed in the same tube the next day (24 hours after its introduction to the arena). In 27 choice tests (out of 45), no decision was made because the choosing queen did not settle in one of the two tubes, or because one of the choice queens escaped their metal wire. Thus in the remaining 18 tests, the choosing queen made a decision, which provided us with 18 chosen queens and 18 not chosen queens. These queens were then kept individually at 21°C, and monitored once a week to count the number of eggs, larvae and pupae present in their tube. The monitoring stopped when the first worker emerged or 80 days after the choice test (12 queens did not produce workers by that time). We used a Leica stereomicroscope to measure the thorax length of the queens at the end of the experiment. Two queens died during monitoring, and were removed from the analysis, as well as the other queen from the same test arena, resulting in 16 chosen and 16 not chosen queens in the analysis.

### Statistics

To investigate the effect of the number of queens on absolute and per capita offspring production, we have extracted for each colony the highest number of brood pieces recorded for each brood type, as well as the number of workers at the end of the experiment, and compared those values between treatment using type II ANOVAs on linear models. To investigate the effect of treatment on the timing of offspring production and development, we compared the days when the first egg, larva, pupa and worker were recorded between colonies with one and two queens using non-parametric Wilcoxon tests. To further investigate the effect of treatment on worker production, we focused on the days after the first workers were produced, and conducted a type II ANOVA on a mixed-effect linear model with the number of workers as response variable, time and number of queens as fixed response variable, and queen identity as a random effect. To investigate the association between size and fecundity in *L. niger* founding queens, we conducted a type II ANOVA on a linear model that explained the number of brood pieces with thorax length (values were standardized within queen collection cohorts for both measurements). To test whether size and fecundity determined queen survival, we compared proportion of surviving queens to 50% using exact binomial tests. To investigate whether chosen and not chosen queens differed in brood production, we extracted for each queen the maximum number of brood pieces recorded for each brood type, and conducted a type II ANOVA on a mixed-effect linear model with the maximum number of brood pieces as response variable, queen category as a fixed response variable, and test arena as a random effect. We performed the same analysis to test the effect of queen category on thorax length. All linear models built in this study were checked for normal distribution of the residuals.

## Results

### Effect of pleometrosis on offspring production

Our comparisons of the number of eggs, larvae, pupae and workers produced in colonies with one or two queens revealed benefits of pleometrosis (Figure 1A). We found that it increased the maximum number of eggs (ANOVA: *F*_1_, _25_ = 39.18, *P* < 0.00001; Figure 1B) and pupae (ANOVA: *F*_1_, _25_ = 6.93, *P* = 0.014; Figure 1D) recorded in the colonies, as well as the number of workers produced (ANOVA: *F*_1_, _20_ = 16.62, *P* = 0.0006; Figure 1E). However, our analysis did not detect such an effect for the maximum number of larvae (ANOVA: *F*_1_, _25_ = 2.25, *P* = 0.15; Figure 1C).

**Figure 1.**
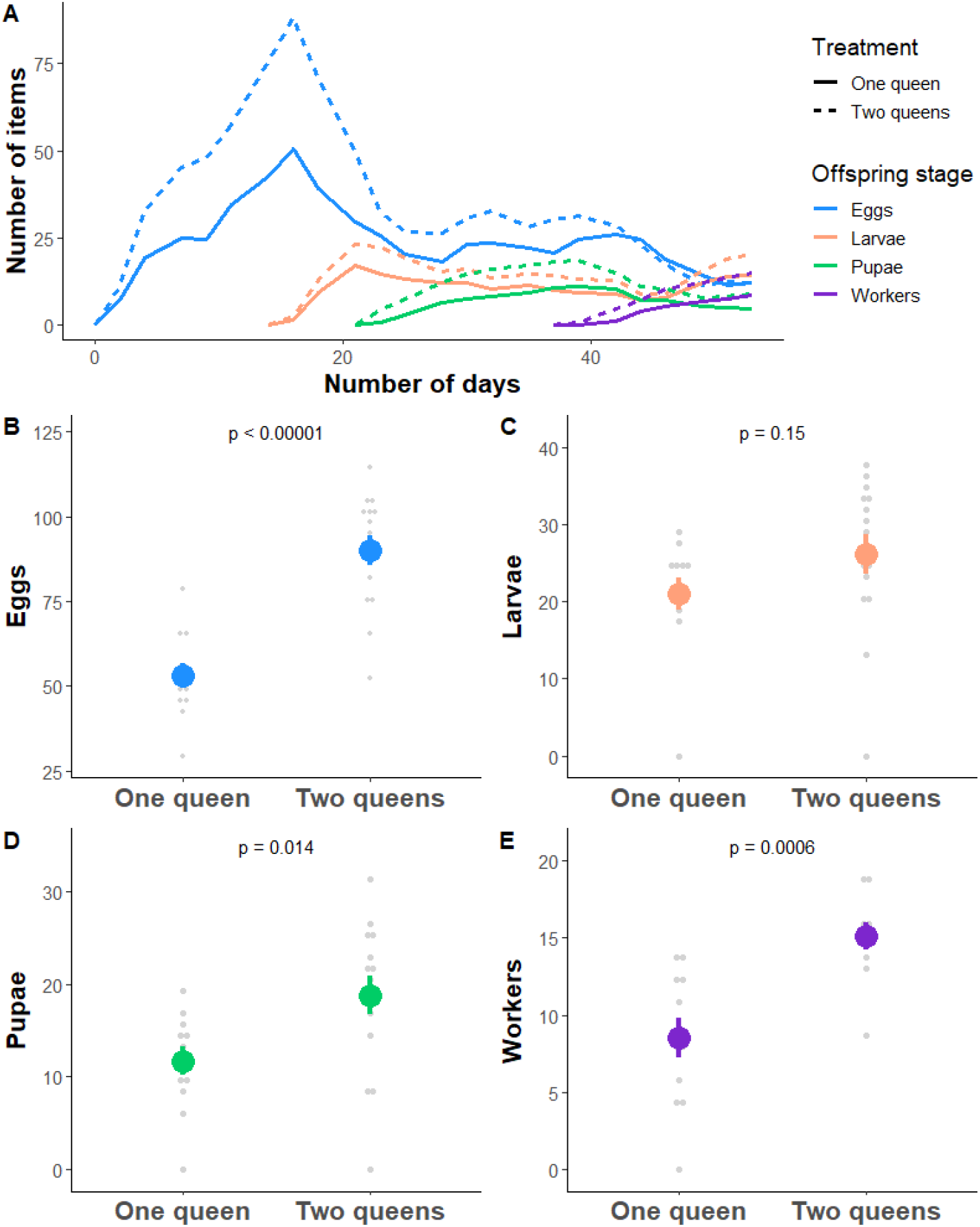
Effect of queen number on A/ mean offspring number over time since colony foundation (colors denote the brood type, while line type indicates treatment), maximum number of B/ eggs, C/ larvae, D/ pupae recorded in the colonies, and E/ number of workers at the end of the experiment (mean ± se).

The benefits of pleometrosis disappeared when correcting for the number of queens present in the colonies. The *per capita* numbers of eggs and larvae were actually reduced in colonies with two queens (ANOVA for eggs: *F*_1_, _25_ = 3.97, *P* = 0.057; ANOVA for larvae: *F*_1_, _25_ = 10.94, *P* = 0.0029), and we found no effect of pleometrosis on the numbers of pupae and workers per queen (ANOVA for pupae: *F*_1_, _25_ = 1.60, *P* = 0.22; ANOVA for workers: *F*_1_, _20_ = 0.43, *P* = 0.52).

We did not detect any effect of queen number on the time to produce the first egg (Wilcoxon test: *W* = 107, *P* = 0.19; Figure 2A) and larva (Wilcoxon test: *W* = 103, *P* = 0.11; Figure 2B), but colonies with two queens were significantly faster than those with one queen in producing their first pupa (Wilcoxon test: *W* = 114.5, *P* = 0.01; Figure 2C) and worker (Wilcoxon test: *W* = 111, *P* = 0.011; Figure 2D).

**Figure 2.**
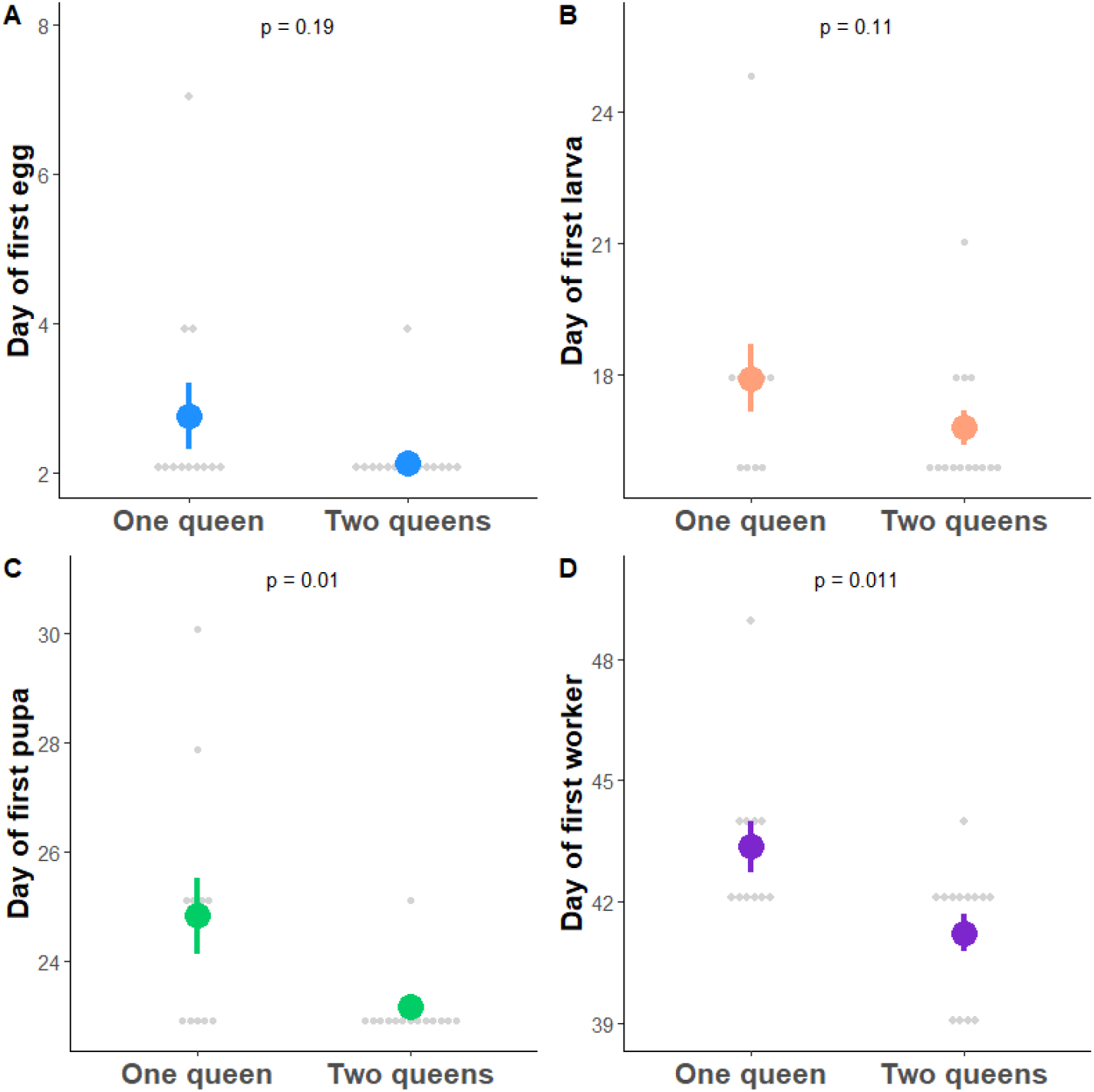
Effect of queen number on the day (mean ± se) when the first A/ eggs, B/ larvae, C/ pupae, and D/ workers were recorded in the colonies.

Finally, once the first workers emerged, colonies with two queens showed a faster increase in worker number than colonies with a single queen (ANOVA: *F*_1,171.4_ = 44.6, *P* < 0.00001; Figure 3).

**Figure 3.**
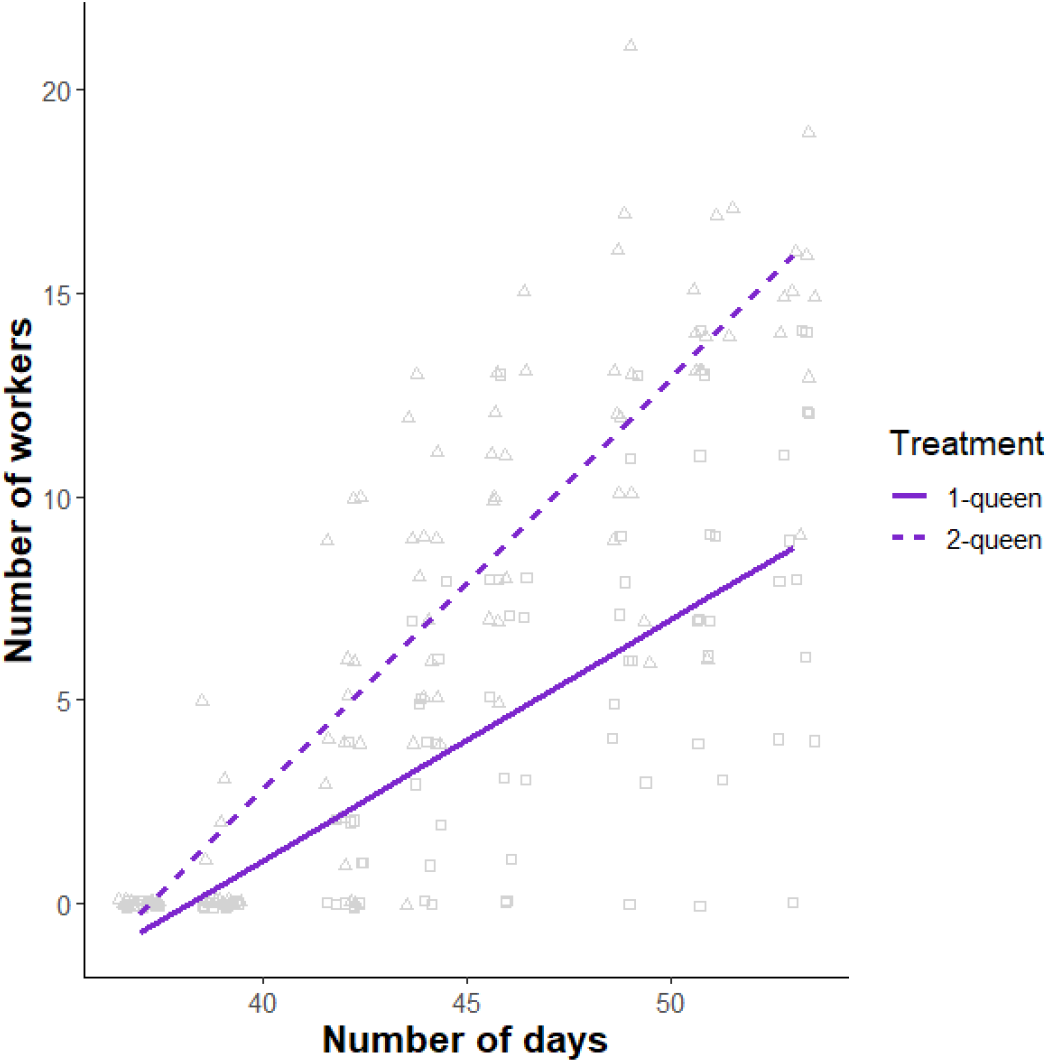
The number of workers increased faster in colonies with two queens (triangle, dotted regression line) compared to those with one queen (square, solid regression line) (ANOVA: F_1,171.4_ = 44.6, P < 0.00001).

### Association between fecundity and size

We found a weak but significant, positive correlation between thorax length and brood number (ANOVA: adjusted *R*² = 0.034, *F*_1_, _229_ = 9.13, *P* = 0.0028). This result confirms that the effects of size and fecundity can be confounded, but suggests that it is possible to disentangle them experimentally.

### Effect of fecundity and size on queen survival

Among the 35 pairs of queens that differed in fecundity (but not size), seven pairs still had two queens at the end of the experiment. Out of the remaining 28 pairs that lost a queen, the most fecund queen survived in 75% (21 out of 28) of the cases, which represented a significant departure from 50% (exact binomial test, *P* = 0.012; Figure 4A). Among the 33 pairs of queens that differed in size (but not fecundity), five pairs still had two queens at the end of the experiment, and in one additional case, the two queens died on the same day. Out of the remaining 27 pairs that lost a queen, the largest queen survived in 63% (17 out of 27) of the cases, which did not differ significantly from 50% (exact binomial test, *P* = 0.25; Figure 4B).

**Figure 4.**
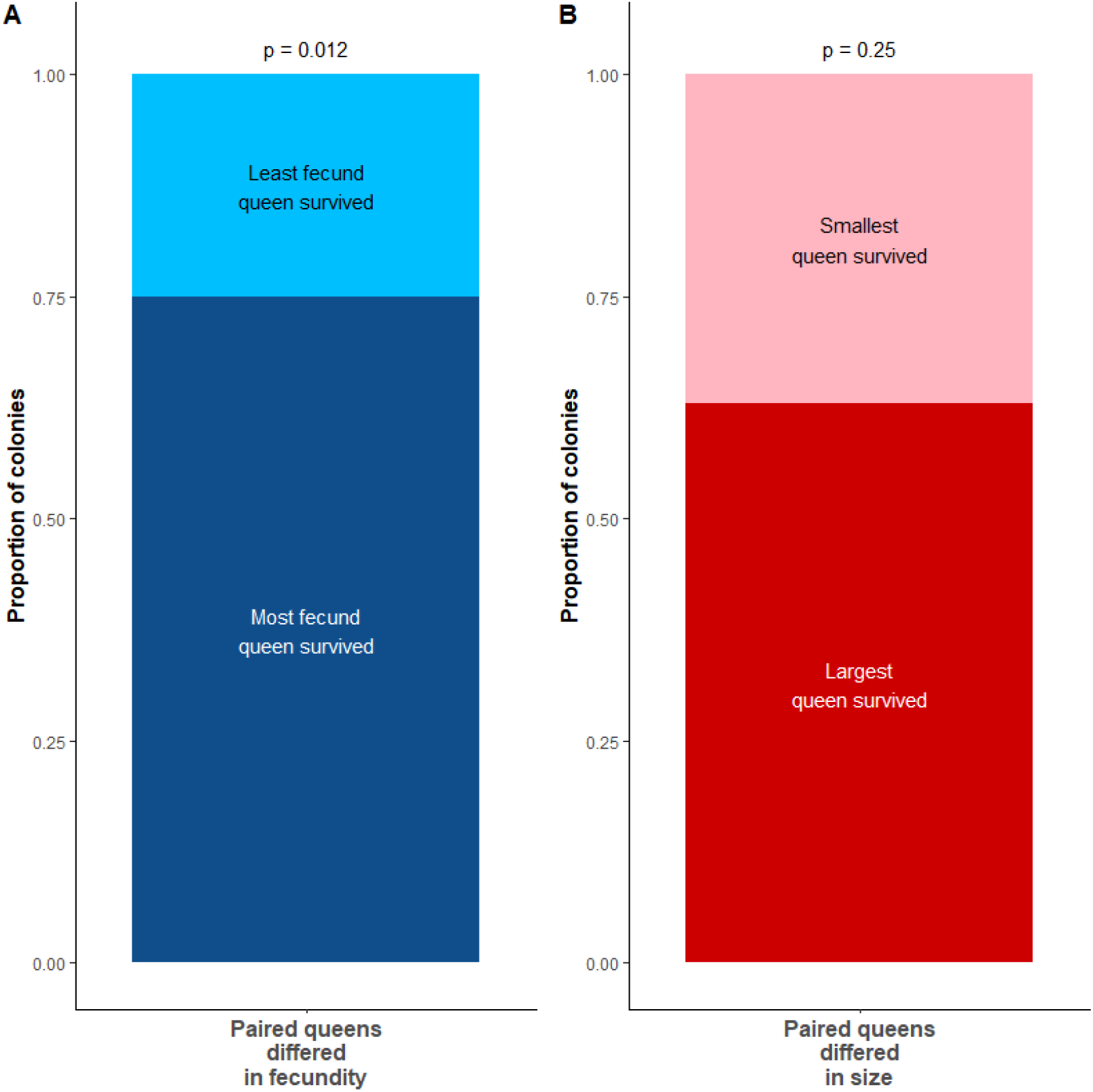
Likelihood of surviving the pleometrotic association depending on queen A/ fecundity and B/ size (p values refer to the comparison to 50%).

### Difference between chosen and not chosen queens in fecundity and size

We found that the maximum number of eggs recorded in the tube was lower for queens that were previously chosen as pleometrotic partners compared to queens that were not chosen (ANOVA: *χ^2^* = 4.56, *P* = 0.033; Figure 5A). We could not find such difference for the maximum number of larvae (ANOVA: *χ^2^* = 0.41, *P* = 0.52) and pupae (ANOVA: *χ^2^* = 0, *P* = 1). In addition, we could not detect any difference in thorax length between chosen and not chosen queens (ANOVA: *χ^2^* = 0.66, *P* = 0.41; Figure 5B).

**Figure 5.**
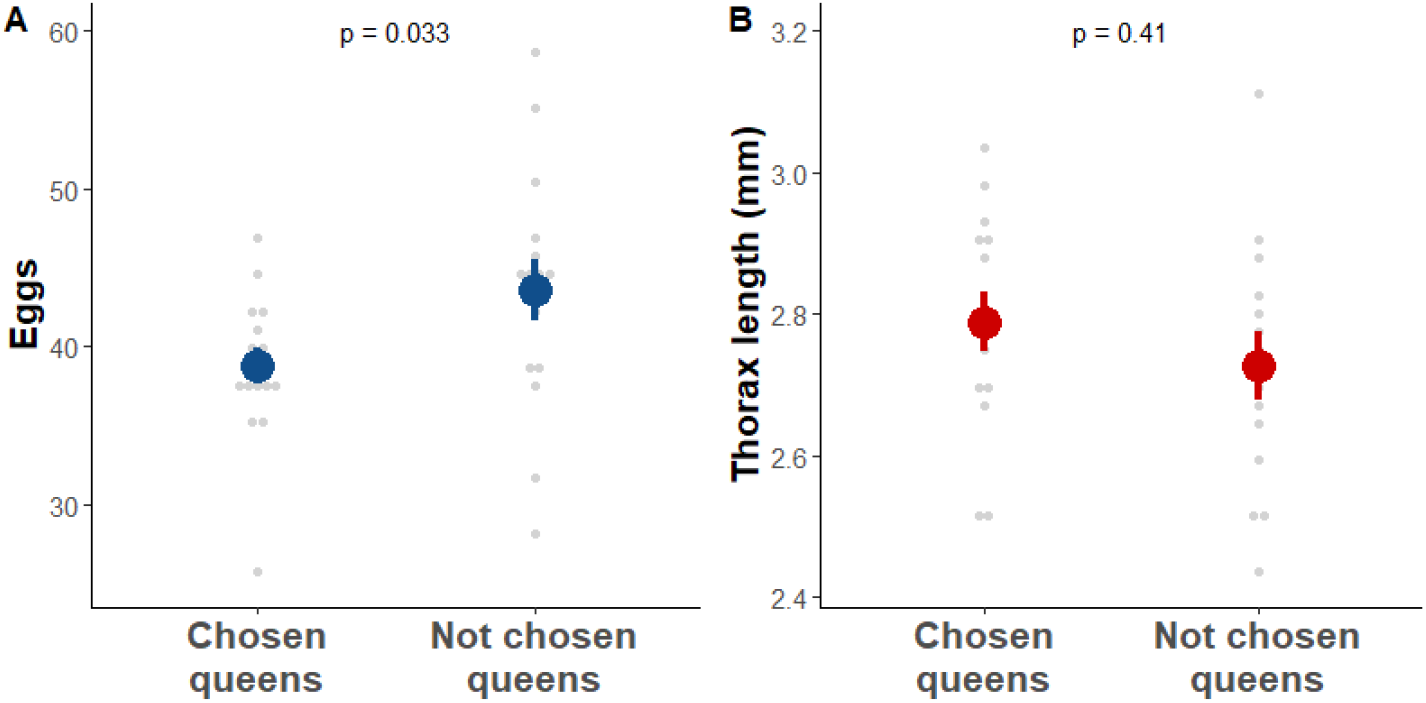
Difference between queens that were chosen and not chosen in the choice tests in A/ maximum number of eggs recorded and B/ thorax length (mean ± se).

## Discussion

In this study, we used the black garden ant *Lasius niger* to investigate the benefits and factors of pleometrosis, the transitory association between founding queens. The monitoring of colonies founded by one or two queens showed that pleometrosis increased and accelerated offspring production. Then, the experimental pairing of *L. niger* founding queens revealed that in pairs of queens of different fecundity but similar size, the most fecund queen was more likely to survive. We did not detect a similar effect of size when controlling for fecundity. Finally, we found that queens associated preferentially with less fecund queens.

Our findings of pleometrosis benefiting offspring production are in line with the literature for this, and other ant species (Clark & Fewell, 2014; Deslippe & Savolainen, 1995; Johnson, 2004; Madsen & Offenberg, 2017; Sommer & Hölldobler, 1995; Trunzer et al., 1998; Waloff, 1957). Interestingly, we only detected these benefits at the colony level, as pleometrosis had either no effect or a negative influence on the *per capita* offspring production. However, colony-level measurements are more relevant in the case of pleometrosis, as the queen that survives the association inherits all the offspring produced during colony foundation. In the field, colonies with a faster, more efficient worker production would have a competitive advantage over neighbouring founding colonies. This is especially true for *L. niger*, which shows high density of founding colonies that compete for limiting resources and raid the brood of other colonies (Madsen & Offenberg, 2017). Thus, the competitive advantage provided by pleometrosis likely enhances colony growth and survival.

The increased and faster production of workers in colonies with two queens may stem from a nutritional boost for the larvae. *L. niger* founding queens do not forage, and produce the first cohort of workers from their own resources. Larvae have been observed to cannibalize both viable and non-viable (trophic) eggs (Urbani, 1991). We found that colonies with two queens produced more eggs, but that this did not translate in them having more larvae. However, more of these larvae became pupae - and ultimately workers. In addition, while the time to produce the first egg and larva did not differ between colonies with one and two queens, the first pupa and worker were produced faster when two queens were present, consistent with a shorter larval stage. We propose that larvae in pleometrotic colonies developed faster and were more likely to reach pupation because they had more eggs at their disposal during development.

These benefits of pleometrosis are only inherited by the queens that survive, it is thus important to understand the factors that determine queen survival in pleometrotic associations. Although this question has been relatively well studied (Aron et al., 2009; Balas & Adams, 1996; Bernasconi & Keller, 1996; Bernasconi & Keller, 1998; Holman et al., 2010; Nonacs, 1990; Sommer & Hölldobler, 1995), it has remained challenging to disentangle the effects of correlated factors. For example, we found that size, which has been reported to predict queen survival (Aron et al., 2009; Bernasconi & Keller, 1998), correlated with fecundity, which would itself be confounded with the parentage of workers in the first cohort produced. To address this issue, we disentangled size and fecundity experimentally, and used workers unrelated to both queens to prevent any potential nepotistic behaviour.

We found that fecundity, but not size, determined queen survival. The finding that, despite being of similar size, more fecund queens are more likely to survive indicates that the outcome of pleometrosis is not the mere consequence of physical dominance. The higher fecundity could reflect a better health condition, which may give the advantage to the more fecund queen in direct fights (Nonacs, 1992), or if workers initiate the fights. Natural selection may have favoured workers that skew aggression toward the less fecund queen, both because this queen would be less efficient at building a colony, and because the workers would be more likely to be the offspring of the more fecund queen. The latter would not necessarily involve direct nepotistic behaviours (the workers would not behave according to parentage, but to fecundity), which have remained elusive in social insects in general (Holzer et al., 2006; Keller, 1997; Ratnieks et al., 2006), and in pleometrotic associations in particular (Balas & Adams, 1996; Bernasconi & Keller, 1996). Despite regular behavioural observations, we did not observe who initiated aggression in our experiments, and it remains unclear whether the queens and/or the workers are responsible for the onset of fights. Consistently with previous studies (Aron et al., 2009; Waloff, 1957), we found that a certain proportion of queen death occurred before worker emergence, suggesting that worker presence is not required for queen execution. Finally, we cannot rule out that the least fecund queens were more likely to die because of a weaker health status, possibly combined with the stress of being associated with another, healthier queen.

Although it has not been directly reported before, our finding that fecundity determines queen survival is consistent with previous reports of weight being associated with queen survival (Bernasconi & Keller, 1996), more fecund queens being more aggressive (Berthelot et al., 2017), CHC profiles differing between winners and losers (Holman et al., 2010), and between more and less fecund queens (Berthelot et al., 2017). We could not directly support previous reports of size correlating with survival (Aron et al., 2009; Bernasconi & Keller, 1998). This could be because in those studies, size could have been confounded with fecundity, and/or because we lacked the statistical power to detect such effect in our experiment.

Pleometrosis provides clear benefits, but these benefits are only inherited by the surviving queens, and the losing queens pay the great cost of dying without contributing to the next generation. Natural selection should thus favour queens that decide whether or not to join a pleometrotic association based on the relative benefits compared to individual foundation - these may differ across ecological contexts (Haney & Fewell, 2018) - and the likelihood of surviving the association. As fecundity appears to determine queen survival in *L. niger*, queens may have evolved the ability to assess the fecundity of potential partners. Our results are consistent with this hypothesis, as queens preferentially associated with queens that would later produce less eggs, possibly because they were less fecund. This further supports our finding that fecundity plays an important role in pleometrotic associations. It is important to note that this difference in egg production could also be a consequence, rather than a cause, of the outcome of the choice experiment. We cannot rule out that entering an association with another queen and/or leaving this association prematurely at the end of the choice experiment may have been stressful for the chosen queen, and affected their later production of eggs. We could not detect any difference between chose and not chosen queens in the number of larvae and pupae produced, which are likely influenced by factors other than fecundity (e.g., brood care behaviour). Interestingly, we did not find that queens chose according to size, consistently with our finding that size may not affect which queen survives the pleometrotic association.

Our study informs on the benefits and factors of pleometrosis, and highlights the role of fecundity in the decision to associate with another queen, and in determining which queen survives the association. It contributes to a better understanding of the onset and outcome of pleometrosis, a classic case of cooperation between unrelated animals.

## Acknowledgements

We thank former and current members of the Behavioural Ecology and Social Evolution group at the JGU Mainz for help with the collection of founding queens, and fruitful discussions on the project.

## Data accessibility

All data is accessible as supplementary material.

## Author contributions

Conceptualization: RL; Methodology and investigation: ET, FD and RL; Analysis, writing, resources and supervision: RL.

